# Skeletal Phenotype and Mechanisms of Bone Loss in *Winnie* Mice as a Model for Inflammatory Bowel Disease

**DOI:** 10.1101/2020.09.28.317495

**Authors:** Ahmed Al Saedi, Shilpa Sharma, Ebrahim Bani Hassan, Lulu Chen, Ali Ghasem-Zadeh, Majid Hassanzadeganroudsari, Jonathan H Gooi, Rhian Stavely, Rajaraman Eri, Dengshun Miao, Kulmira Nurgali, Gustavo Duque

## Abstract

**Objective:** We aimed to investigate the skeletal phenotype of *Winnie* mouse model of spontaneous chronic colitis, which carries a mutation in the *Muc2* gene and closely replicates IBD symptoms and pathophysiology. These mice have a high level of gut-derived serotonin (GDS), a potent osteoblastogenesis inhibitor. We explored the underlying mechanisms of bone loss associated with chronic intestinal inflammation.

**Design:** *Winnie* male and female mice prior to colitis onset (6 weeks old) and progression (14 and 24 weeks) were compared to age- and sex-matched C57BL/6 controls. We assessed bone quality (static and dynamic histomorphometry, micro-CT, 3-point bending), intestinal inflammation (lipocalin-2), GDS levels, serum levels of calcium, phosphorus and vitamin D, *ex vivo* bone marrow analysis and molecular mechanisms inhibiting osteoblastogenesis.

**Results:** Significant deterioration in trabecular and cortical microarchitecture, reductions in bone formation, mineral apposition rate, bone volume, osteoid volume and bone strength were observed in *Winnie* mice compared to C57BL/6 controls. Decreased osteoblast and increased osteoclast numbers were prominent in *Winnie* mice. We report for the first time that elevated GDS cross-talks with molecular pathways to inhibit bone formation in *Winnie* mice. Increased expression of 5-HTR1B and FOXO1 mRNAs, dissociation of FOXO1/CREB1 complex and association of FOXO1 with ATF4, promoting the transcriptional activity of FOXO1, results in suppression of osteoblast proliferation in *Winnie* mice compared to controls.

**Conclusion:** These findings open avenues for the development of targeted therapies for IBD-related bone loss.

**Significance of this study:** *What is already known on this subject?:*

- Osteoporosis is a common extraintestinal manifestation of inflammatory bowel disease (IBD).
- Currently available treatments are not effective for IBD-associated bone loss.
- The mechanisms of bone loss are poorly understood. A major limitation has been the lack of an appropriate animal model for IBD-associated bone loss.

*What are the new findings?:*

- We report for the first-time the skeletal phenotype in *Winnie* mouse model of IBD
- This study presents a novel mechanism of IBD-associated bone loss, involving elevated gut-derived serotonin crosstalk with molecular pathways inhibiting bone formation.

*How might it impact on clinical practice in the foreseeable future:*

- These findings open avenues for the development of targeted therapies for IBD-related bone loss.

## Introduction

Inflammatory bowel diseases (IBD) are chronic, relapsing and progressively debilitating disorders, including Crohn’s disease (CD) and ulcerative colitis (UC).^1^ Genetic susceptibility, immune system dysfunction, environmental factors and shifts in gut microbiota have been implicated with IBD development.^2,3^ IBD is a systemic disorder that affects many organs with musculoskeletal manifestations being the most common.^4^ It is estimated that 14–42% of IBD patients have osteoporosis and therefore, a considerably high incidence of fragility fractures (40%).^5^ This results in increased disability, reduced quality of life and risk of mortality, especially amongst older patients with spine and hip fractures.^6,7^Chronic inflammation, mainly mediated by tumor necrosis factor (TNF)-α, leads to bone loss via activation of the receptor activator of the nuclear factor (NF)-κB (RANK) ligand RANKL/RANK/osteoprotegerin (OPG) system, which promotes osteoclastogenesis.^8^ Other factors, such as malabsorption of essential nutrients like calcium and vitamin D and longterm use of glucocorticoids (GCs), contribute to bone loss in IBD patients.^9,10^ Despite significant advances in uncovering the pathogenesis of IBD, the precise mechanisms underlying IBD-associated bone loss have not been fully elucidated.

Mouse models have been widely used to understand pathophysiological mechanisms of IBD and IBD-associated co-morbidities. Skeletal deterioration has been previously reported in chemically-induced animal models of colitis (e.g. induced by 2,4,6-trinitrobenzene sulfonic acid (TNBS) or dextran sulfate sodium (DSS)).^11,12^ Drastic reductions in trabecular and cortical bone volumes were noticed during the acute phase of DSS-induced colitis. However, bone measurements restored to control when pro-inflammatory cytokines returned to normal levels.^12^ In a chronic model of DSS colitis, increase in bone TNF-α levels negatively correlated with bone mass, and was independent of weight loss.^13^ While TNBS or DSS-induced colitis models provided valuable information about bone loss, but inflammation lasted only 7–15 days after the induction. Therefore, the mechanisms involved in the longterm association between chronic inflammation and skeletal health deterioration could not be validated in these models. In addition, correlation between the severity of colitis and bone loss was observed in a chronic colitis model, *Helicobacter hepaticus*-infected interleukin-10-deficient male mice.^14^ However, colitis in this model only could be studied in male mice as *Helicobacter hepaticus* survival and virulence is gender-dependent.^14^

*Winnie* mouse model of spontaneous chronic colitis corroborates as an *in vivo* and *ex vivo* human model system to study IBD.^15^ *Winnie* mice possess a missense mutation in the *Muc2* gene resulting in aberrant Muc2 mucin biosynthesis, reduction of mucin storage in goblet cells, and accumulation of non-glycosylated Muc2 precursor within goblet cells.^15,16^ In humans, reduced levels or absence of *Muc2* gene expression occurs in CD, while in active UC, Muc2 production and secretion are reduced, leading to a thinner mucosal layer and increased intestinal permeability.^17,18^ The primary epithelial defect and mucosal barrier dysfunction in *Winnie* mice results in complex multi-cytokine mediated colitis involving both innate and adaptive immune components with a dominant IL-23/TH17 response similar to IBD patients.^15^ The similarities between *Winnie* mice and UC patients have been further confirmed by determining alterations in colon morphology, changes to colonic motility, transit time, fecal microbial and metabolomic profiles.^19–21^

Mechanisms involved in IBD-associated bone pathology are not yet clear. Recent evidence implicates the role of tryptophan (TRP) and TRP metabolites in bone homeostasis and loss.^22^ One of the TRP metabolites, picolinic acid, has an anabolic effect on bone^23^, while other TRP products have a contrary action on bone e.g. gut-derived serotonin (GDS), which can decrease bone mass via activation of receptors on pre-osteoblasts.^24^ To date, only one study has reported a potential link between high GDS levels and bone deficits in DSS model.^25^ However, in this study GDS-mediated effects on bone pathology have been revealed for 21 days of colitis only. Interestingly, a high level of GDS is prominent in *Winnie* mice.^26^ Unlike studies in models of chemically-induced colitis, *Winnie* model permits long-term bone studies throughout the chronic disease course up to ~6-7 months mimicking the onset and progression of UC.

In this study, we aimed to characterize the skeletal phenotype of *Winnie* mice compared to age and sex-matched control C57BL/6 mice and to investigate mechanisms of bone pathology by testing the role of increased GDS levels in the downstream bone formation pathways.

## Materials and Methods

### Animals

Male and female *Winnie* mice (C57BL/6 background) were obtained from the University of Tasmania (Launceston, Australia) (animal ethics: AEC17/016). Age- and sex-matched control C57BL/6 mice were obtained from the Animal Resource Centre (Perth, Western Australia). All mice were acclimatized for one week in a temperature-controlled animal holding facility with a 12hr day/night cycle and *ad libitum* access to food/water. Mice were euthanized by a lethal dose of pentobarbital at 6, 14 and 24wk of age and tibiae and femora bones, blood and gastrointestinal tissues were collected.

### Micro-computed tomography (Micro-CT)

Femora bones fixed with 4% paraformaldehyde (PFA) underwent Micro-CT scanning (SkyScan 1272, Bruker, Belgium) with isotropic voxel size (resolution) of 5.4μm, (n=4-8/group). All images were reconstructed and analyzed using manufacturers’ guidelines. The reported bone microstructure parameters are based on ASBMR guidelines.^27^

### Three-point bending test

Mechanical testing of femora from C57BL/6 and *Winnie* mice (n=4-8/group) was performed by three-point bending tests as previously described.^28^ All mechanical testing was conducted with the observer being blinded to the genotype of the samples.

### Histology and histochemical analysis

Following PFA fixation of tibiae from controls and *Winnie* mice from all 3 age groups (n=4-8/group) for 24hrs, bones were decalcified in 14% EDTA (pH 7.4) for 2wk. Bones were embedded in paraffin and 5μm sections were cut using a rotary microtome (Leica RM2255, Germany). Sections were stained with hematoxylin and eosin (H&E) for osteoblasts and tartrate-resistant acid phosphatase (TRAP) for osteoclasts as previously described.^29^

### Total collagen staining

Tissue sections were deparaffinized, hydrated through xylenes and graded alcohol series and mounted with Kaiser’s glycerol jelly (n=4-8/group) followed by incubation in picronitric acid-direct red solution for 0.5-1hr at room temperature (RT), washing, counterstaining with hematoxylin and rinsing in running water.^29^

### Immunostaining for type I collagen

Sections were de-waxed and rehydrated with phosphate-buffered saline (PBS) (n=4-8/group), incubated with goat anti-type 1 collagen antibody (Ab) (1:100, ab34710, Abcam, UK) overnight at RT, washed, incubated with rabbit anti-goat IgG (1:100, BD349031, BD Biosciences, NJ, US) at 28°C for 1hr and washed. Sections were stained with Elite-ABC at 28°C for 1hr (Vector Elite ABC kit).^29^ Finally, staining was done using Diaminobenzidine (DAB) followed by counterstaining with haematoxylin.

### Plastic-embedded sectioning

Tibiae bones were incubated with 100, 95, 90 and 70% ethanol for 24hrs each at 4°C (n=4-8/group). Bones were rinsed with PBS and embedded in polymethyl methacrylate (MMA). Serial 4-6μm sections of MMA-embedded tissues were stained with Von Kossa histochemical staining for bone matrix. The primary histomorphometric data were obtained using Bioquant^®^ image analysis software (OSTEO 19.6.6, US).

### Dynamic histomorphometry by calcein dual-labelling experiment

The fluorochrome calcein (10mg/kg dissolved in 2.0% sodium bicarbonate, pH 7) was injected at 0.1mL/10g body weight 7 and 2 days prior to harvesting bones from mice. The tibia condyles were dissected and embedding, and ultra-thin sectioning were done in a dark room (n=4-8/group). Fresh tissue sections were observed and imaged under a ZEISS AxioImager2 fluorescence microscope, equipped with a Retiga 1300 camera (ZEISS^®^, Germany).

### Ex vivo bone marrow analysis

Bone marrow from *Winnie* and control mice tibiae (n=4-8 mice/group) was collected by flushing with α-modified essential medium ((α-V1EV1, Sigma, Australia). Bone marrow-derived mesenchymal stromal cells (BM-MSCs) were isolated by their adherence to plastic and cultured at a second passage. At day 7 of culture, osteoblast differentiation capacity was assessed by alkaline phosphatase (ALP) staining using SIGMAFAST™ p-Nitrophenyl phosphate tablets (N1891, Sigma). Colonies with more than 10% of ALPpositive cells were considered as colony-forming units–osteoblasts (CFU-OBs). Matrix mineralization was stained and quantified by extracting Alizarin red with 100mM cetylpyridinium chloride at RT at day 14. The absorbance was measured at 562nm spectrophotometrically.

### Serum analysis for calcium, phosphorus, and vitamin D

Calcium, phosphorus, and vitamin D levels were assayed by ASAP Laboratory services (Melbourne, Australia) according to the manufacturer’s instructions (n=4-6/group). Calcium ions reacted precisely with Arsenazo III to form purple-colored calcium-arsenazo III complex. Absorbance was measured at 660/700nm. Inorganic phosphate reacted with molybdate to form a heteropolyacid complex. Absorbance was measured at 340/380nm, while vitamin D levels were measured by radioimmunoassay (Diagnostic Products).

### Quantitative Real-time PCR

Purified total RNA was extracted from fresh BM-MSCs at passage 2 using a Qiagen Minikit (Qiagen, Australia). Quantitative RT-PCR was performed (n=3 mice/group, 2 repeats) as previously described.^26^ The mRNA levels of HTR1B, FOXO1, and reference gene GAPDH were detected using PrimePCR™ Assays (Bio-Rad, Australia). The mRNA level of each gene was normalized to GAPDH and expressed relative to C57BL/6 mice using the 2-ΛΛCt method.

### Assessment of gut-derived serotonin (GDS)

Electrochemical measurements of GDS at the mucosal surface of the colon was performed using a carbon fiber electrode to mechanically stimulate and oxidize GDS to generate a compression-evoked (peak) release of GDS which decays back to basal levels (steady state) in both control and *Winnie* mice (n=4-6/group) as previously described.^26^

### Fecal analysis for Lipocalin-2 protein

Quantitative measurement of fecal lipocalin-2 (neutrophil gelatinase-associated lipocalin, NGAL) was performed using a SimpleStep ELISA^®^ kit according to the manufacturer’s instructions (ab119601) (n=7-11/group). This method for non-invasive measurement of intestinal inflammation is routinely used in our lab.^19^

### Double immunofluorescence analysis

Double immunofluorescence labeling was performed in BM-MSCs (n=3-4 mice/group). BM-MSCs at passage 2 were fixed in 4% PFA for 20min at RT and incubated with mouse monoclonal FOXO1 Ab (14952, Cell Signalling Technology, MA, US), rabbit polyclonal anti-phospho CREB1 conjugate (06-519-AF647, Merck, Germany) and rabbit polyclonal activating transcription factor-4 (ATF4) (ab23760, Abcam, UK) in Ab dilution buffer (1x PBS/1% BSA/0.3% Triton™ X-100) overnight at 4°C, and washed with PBS. Both types of primary Ab labelled BM-MSCs (CREB1:FOXO1) and (ATF4: FOXO1) were incubated with secondary antibodies, FITC-conjugated anti-mouse IgG (1:250), Cy3-conjugated rabbit antimouse IgG (1:200) for 2hrs at RT in the dark. Labeled BM-MSCs were visualized and imaged with an Eclipse Ti confocal microscope (Nikon, Japan) using 60X objective.

### Statistical analysis

Data were analysed by using Student’s *t* test (two-tailed) and two-way analysis of variance (ANOVA) when appropriate for multiple group comparisons followed by Tukey’s and Sidak’s post hoc tests using GraphPad Prism V.7. A *p* value of <0.05 was considered to indicate statistical significance.

## Results

### Alterations of microstructure and biomechanical competence in the femoral metaphysis of Winnie mice with spontaneous chronic colitis

Bone microstructure, cellular activity and strength were assessed in female and male *Winnie* mice with spontaneous chronic colitis at different age groups based on severity of colitis: 6wk (no colitis), 14wk (progressive colitis) and 24wk (severe colitis). Micro-CT data showed significant deterioration in trabecular and cortical microarchitecture in *Winnie* mice at 14wk and 24wk compared to age-matched controls. Indeed, significant decreases in bone volume to total volume ratio (BV/TV) were observed in female *Winnie* mice (−34% at 14wk, −41.5% at 24wk) (**Figure 1A**) compared to age- and sex-matched controls and to 6wk female *Winnie* mice. While BV/TV in 6wk female *Winnie* was similar to age-matched female controls, the male *Winnie* mice showed lower BV/TV in all 3 age groups. We observed significantly lower trabecular thickness (Tb.Th) in both male and female *Winnie* mice at 24wk compared to age- and sex-matched controls (**Figure 1A, 1B**). No difference was found in trabecular separation (Tb.Sp) in female and male *Winnie* mice at any age. Lower trabecular number (Tb.N) was observed in male *Winnie* mice at 14wk compared to age-matched male controls. However, no detectable changes were seen in Tb.N in female *Winnie* mice at any age. Both female and male *Winnie* mice showed significant reduction in cortical thickness (Ct.Th) at 14wk and 24wk. Both genders of *Winnie* and control mice at 6wk showed similar Ct.Th.

**Figure 1:**
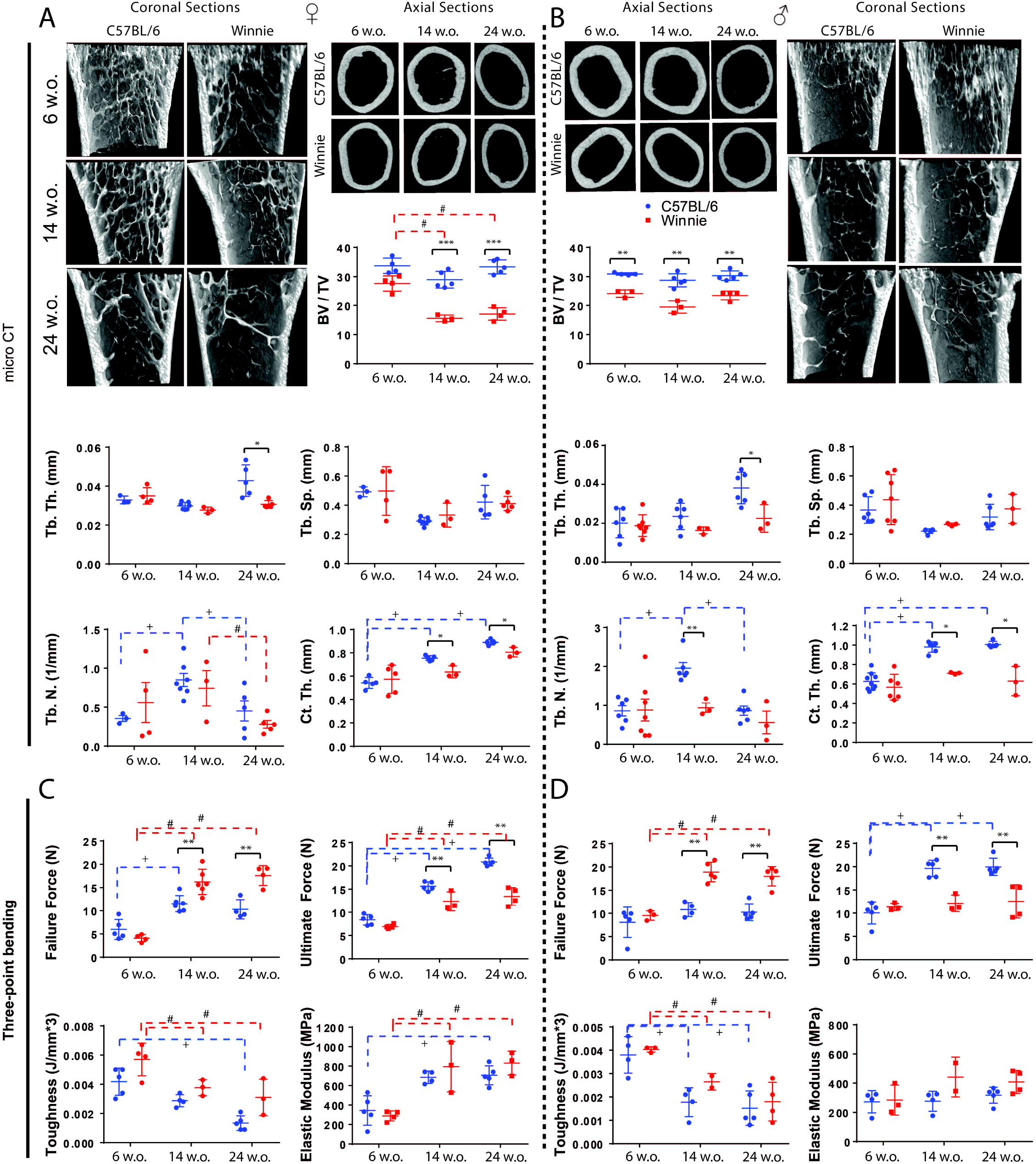
Quantitative analysis of microcomputed tomographic (μCT) images and three-point bending analysis of femoral bones from Winnie mice compared to C57BL/6 controls. **A, B)** Representative longitudinal and cross-sectional images of three-dimensional reconstructed distal ends and diaphysis of femora from female (**A**) and male (**B**) control and *Winnie* mice at 6wk, 14wk and 24wk of age utilizing microcomputed tomography. Parameters analyzed: Bone Volume (BV)/ Total Volume (TV), Trabecular Thickness (Tb.Th), Trabecular Separation (Tb.Sp), Trabecular Number (Tb.N), Cortical Thickness (Ct.Th). **C, D)** Three-point bending analysis of femora from female (**C**) and male (**D**) *Winnie* mice compared to C57BL/6 controls at 6wk, 14wk and 24wk of age. Parameters analysed: Failure Force (N), Ultimate Force (N), Toughness (J/mm*3) and Elastic Modules (E). Each value is the mean±SEM, n=4-8/group. \**p*<.05; \*\**p*<.01; \*\*\**p*<.001 *Winnie* mice compared to C57BL/6. **#***p*<.05 differences between age groups of *Winnie* mice. **+***p*<.05 differences between age groups of C57BL/6 mice.

Furthermore, we investigated the bone mechanical properties by 3-point bending. We found significantly increased failure force (load) (N) in both female and male *Winnie* mice at 14wk and 24wk compared to age- and sex-matched controls. No difference in failure force was observed between genotypes at 6wk. However, females were significantly lower than males in both genotypes (**Figure 1C, 1D**). Ultimate force (N) decreased significantly in both female and male *Winnie* mice at 14wk and 24wk compared to age- and sex-matched controls. Toughness and elastic modulus were not significantly different between *Winnie* and controls at all ages in both genders. While toughness was reduced at 14wk and 24wk compared to 6wk in both genders and genotypes, elastic modulus was increased significantly in female *Winnie* mice at 14wk and 24wk and in female controls at 24wk (**Figure 1C, 1D**).

### Histomorphometric analysis showed reduction in osteoblast numbers and collagen I, with simultaneous increase in osteoclast numbers

The apparent catabolic effect of colitis on bone was further examined by quantifying the number of osteoblasts, collagen, and osteoclasts *in situ*. After normalization to the bone surface (B.S), a significant decrease in osteoblast numbers (Ob/B.S) was observed in all age groups in both genders of *Winnie* mice (**Figure 2A**). In female *Winnie* mice, Ob/B.S were decreased by −61% at 6wk, −41.5% at 14wk and −54% at 24wk compared to controls. In male *Winnie* mice, Ob/B.S were decreased by −41.8% at 6wk, −39.2% at 14wk and −38.8% at 24wk compared to age- and sex-matched controls. Collagen-I was significantly reduced in female *Winnie* mice by −8.2% at 6wk, −52.6% at 14wk, −61.2% at 24wk. Similarly, in male *Winnie* mice collagen-I was reduced by −7.1% at 6wk, −80.8% at 14wk, −78.5% at 24wk compared to male controls (**Figure 2B**). Concomitant to the effect observed in the osteoblast numbers and marked elevation in osteoclast numbers after normalization to the bone surface (Oc/B.S.) was prominent in *Winnie* mice. Increases in Oc/B.S. in female *Winnie* mice were +51% at 14wk and +31% at 24wk compared to age-and sex-matched controls. In male *Winnie* mice, there were increases in Oc/B.S. by +60% at 14wk, +38% at 24wk. No difference in Oc/B.S. was observed at 6wk in both genders (**Figure 2C**).

**Figure 2:**
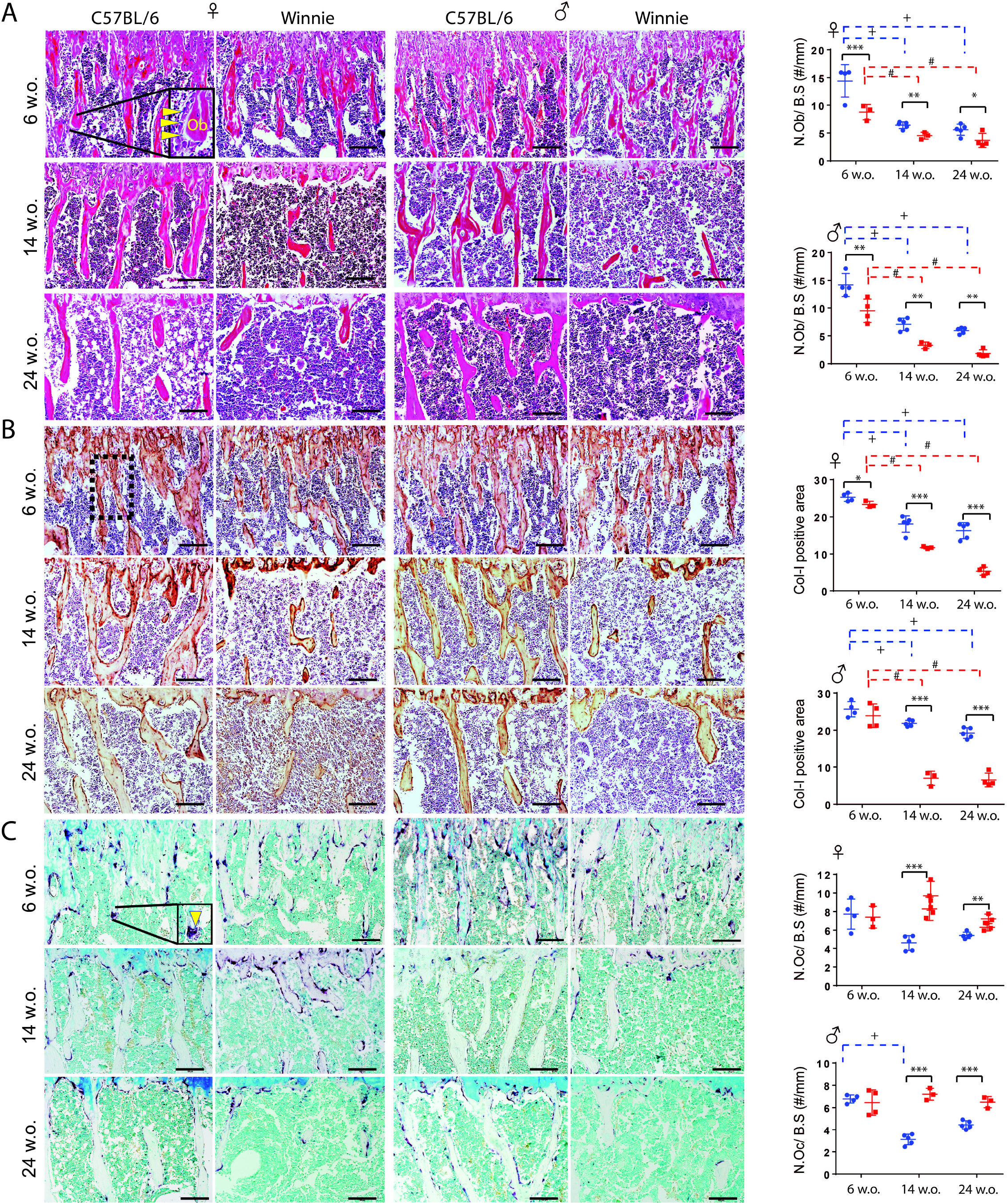
Bone niche cells in Winnie mice compared to C57BL/6 controls. Representative micrographs of paraffin-embedded sections of tibial metaphyseal regions from 6wk, 14wk and 24wk of age C57BL/6 and *Winnie* mice. **A)** Hematoxylin and Eosin staining (H&E) to show active osteoblast, osteoblast numbers (N.Ob) relative to bone surface (B.S) (N.Ob/B.S, #/mm), scale bars represent 100 □μm; **B)** Representative micrographs of tibial sections stained for type I collagen (Col-I) to stain Col-I positive immunoreactive areas, scale bars represent 100μm; **C)** Tartrate-resistant acid phosphatase (TRAP) staining for osteoclast number relative to bone surface (N.Oc/B.S, #/mm), scale bars represent 100μm. Each value is the mean□±□SEM, n=4-8/group. \**p*<.05; \*\**p*<.01; \*\*\**p*<.001 *Winnie* mice compared to C57BL/6. **#***p*<.05 differences between age groups of *Winnie* mice. **+***p*<.05 differences between age groups of C57BL/6 mice.

### Static and dynamic histomorphometry analyses showed low bone formation, mineral apposition rate, and differentiation capacity of BM-MSCs in Winnie mice

Noteworthy reduction in BV/TV (total collagen) was observed in 3 *Winnie* age groups (both females and males) compared to age- and sex-matched controls. BV/TV (T-Col) was reduced in female (−11.5% at 6wk, −58% at 14wk, −61% at 24wk) and male (−10% at 6wk, −47% at 14wk, −63.5% at 24wk) *Winnie* mice compared to age-and sex-matched controls. The ratio of the osteoid volume to the bone volume (OV/BV) was significantly reduced in female (−21.5% at 14wk, −23.2% at 24wk) and male (−22.3% at 14wk, −24.5% at 24wk) *Winnie* mice compared to controls. Although no significant decrease in OV/BV was observed in 6wk *Winnie* mice compared to controls, OV/BV was significantly lower at 14wk and 24wk compared to 6wk *Winnie* mice of both genders.

In addition, quantitative dynamic histomorphometry performed on dual-labelled sections revealed decreased mineral apposition rates (MAR) in trabecular bone in *Winnie mice* compared to controls. *Winnie* mice represented lower MAR in females (−31.6%, −29.5%) and males (−18% and −24%) at 14 and 24wk compared to controls and 6wk *Winnie*. Furthermore, undecalcified histology showed evidence of decreased trabecular bone formation rate (BFR/BV) in *Winnie* females at 14wk (−29%) and 24wk (−36.5%) and males at 14wk (−34%) and 24wk (−35%) (**Figure 3E, 3F**). In both male and female *Winnie* mice, BFR/BV progressively reduced with age. Mineralization capacity was significantly decreased in *Winnie* mice at 14wk and 24wk in both females and males compared to age and sex-matched controls, but not at 6wk (**Figure 3G, 3H**). The number of colony-forming unit osteoblasts (CFU-OBs) were significantly reduced in female and tend to decrease in male *Winnie* mice at 14wk compared to controls, while reduced in both genders at 24wk. No differences in CFU-OBs were observed at 6wk male and female *Winnie* mice compared to age-matched controls (**Figure 3I, 3J)**.

**Figure 3:**
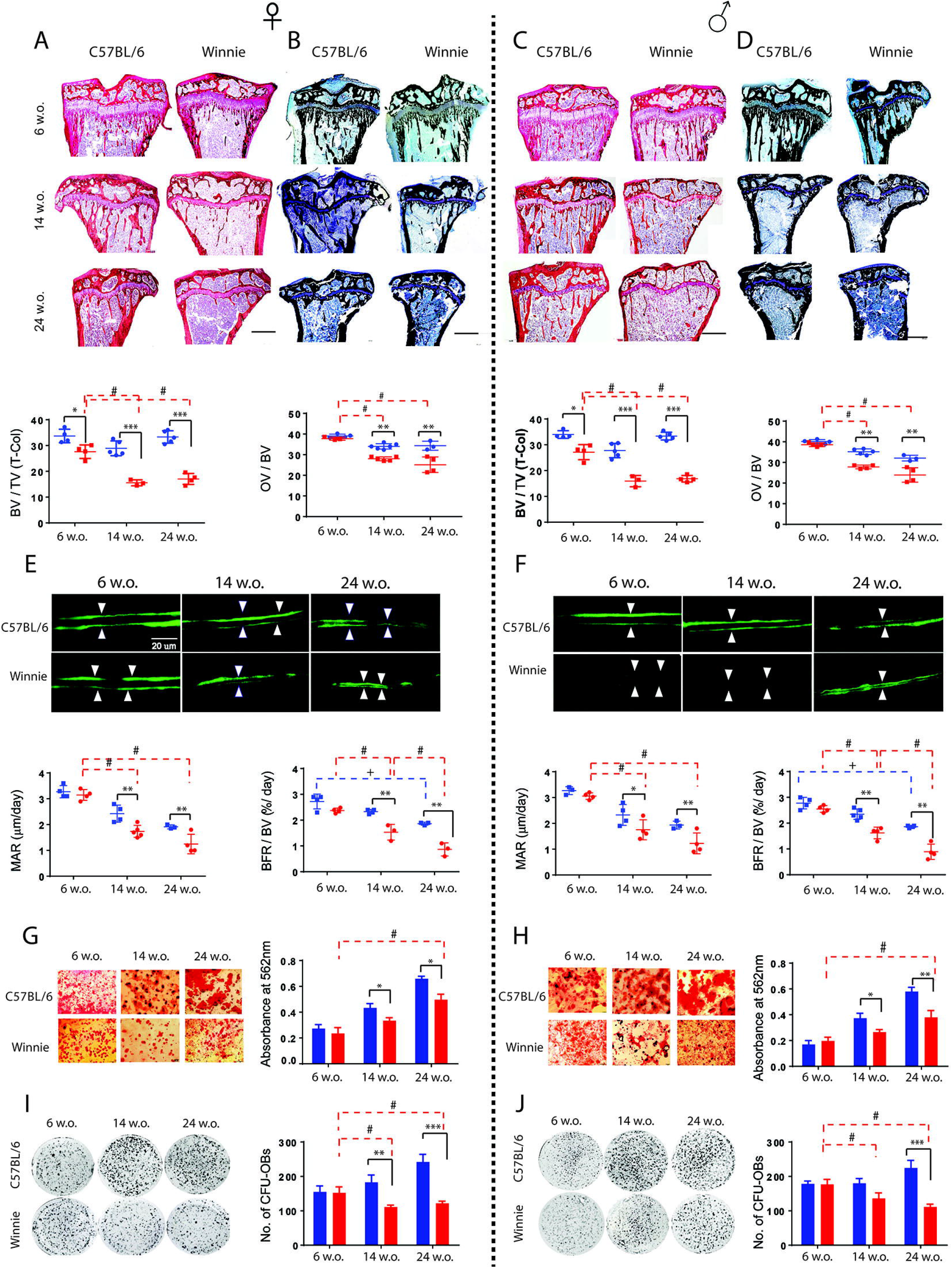
Analysis of bone formation and differentiation of bone marrow mesenchymal stem cells (BM-MSCs) in Winnie mice compared to C57BL/6 controls. **A, C)** Histochemical staining for total collagen of paraffin □ embedded sections of tibiae from female (**A**) and male (**C**) *Winnie* and control C57BL/6 mice at 6wk, 14wk, 24wk of age; analysis of total collagen (T-Col) utilizing Bone Volume/Total volume (BV/TV) ratio. Scale bars represent 500μm. **B, D)** Von Kossa staining of plastic embedded sections of tibiae from female (**B**) and male (**D**) *Winnie* and control C57BL/6 mice at 6, 14, 24 weeks of age; mineralization level is analysed utilizing Osteoid Volume/Bone Volume (OV/BV) ratio. Scale bars represent 500μm. **E, F)** Representative images of bone formation visualized in the femur trabecular bone from female (**E**) and male (**F**) *Winnie* mice at 6wk, 14wk, 24wk of age compared to age-matching control C57BL/6 mice by dual-labelling for Calcium. Scale bars represent 20μm. Bone formation was analyzed utilizing Mineral Apposition Rate (MAR, μm/day) and Bone Formation Rate (BFR)/Bone volume (BV) (%/day). **G, H)** Primary BM-MSCs from *Winnie* and control C57BL/6 mice at 6wk, 14wk, 24wk of age were cultured *ex vivo* in osteogenic differentiation medium for 21 days. Resulting cultures were stained with Alizarin Red to show mineralization of BM-MSCs from female (**G**) and male (**H**) mice (absorbance at 562nm) and alkaline phosphatase (ALP) to show colony-forming unit osteoblasts (CFU-OBs) of BM-MSCs from female (**I**) and male (**J**) mice. Each value is the mean□±□SEM, n=4-8/group. \**p*<.05; \*\**p*<.01; \*\*\**p*<.001 *Winnie* mice compared to C57BL/6. #*p*<.05 differences between age groups of *Winnie* mice. +*p*<.05 differences between age groups of C57BL/6 mice.

### Increased HTR1B and FOXO1 expression in response to elevated GDS levels and its correlation with the changing partners of FOXO1

To investigate relationship between higher GDS levels and bone pathology, we analyzed downstream molecular players of GDS on bone by qRT-PCR. The gene expression analysis of BM-MSCs at passage 2 revealed significantly higher levels of *5-HTR1B* and *FOXO1* genes relative to GAPDH reference genes at 14wk *Winnie* (females and males) compared to age- and sex-matched controls (2-ΔΔCt=1.00) (**Figure 4A, 4B**). However, relative gene expression levels of *5-HTR1B* and *FOXO1* in BM-MSCs of *Winnie* mice at 6wk (females and males) was similar to age and sex-matched controls.

**Figure 4.**
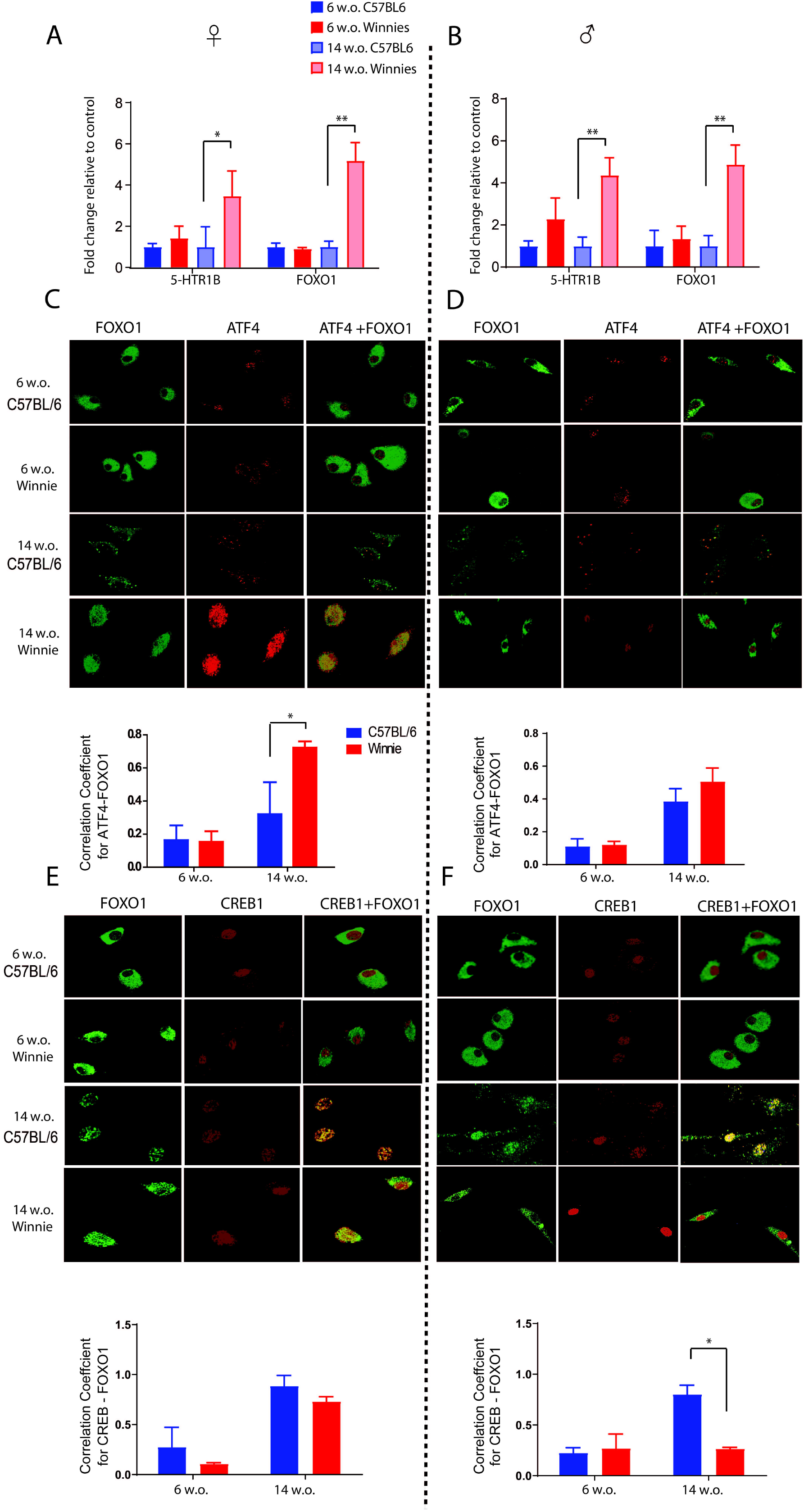
Mechanisms of decreased osteoblastogenesis in Winnie compared to C57BL/6 controls. **A, B)** Quantitative 5-HTR1B and FOXO1 genes expression analysis in bone marrow stromal cells in *Winnie* mice and C57BL/6 controls in 6wk and 14wk females **(A)** and males **(B)** normalized to GAPDH reference gene; n=3 mice/group. **(C, D)** Confocal Microscopy of subcellular compartments for testing colocalization of FOXO1-ATF4 was performed in female BM-MSCs **(C)** and males **(D)**. **(E, F)** Confocal Microscopy of subcellular compartments for testing colocalization of FOXO1-CREB1 was performed in BM-MSCs from female **(E)** and male **(F)** mice. Representative confocal microscopy images at 60X. Scale bar: 50μm, n=3-4 mice/ group. Statistical significance, \**p*<.05 \*\**p*<.01 and \*\*\**p*<.001 vs. control.

In order to find FOXO1 transcriptional partners that differentially regulate its activity in the presence and absence of high GDS levels, we investigated interactions of downstream target molecules, FOXO1 association with activating transcription factor 4 (ATF4) and cAMP-responsive element binding protein 1 (CREB1). BM-MSCs from 14wk female *Winnie* mice showed enhanced association of ATF4 with FOXO1 (ATF4-FOXO1) compared to 14wk female controls (**Figure 4C**), but not in males (**Figure 4D**). ATF4-FOXO1 complex formation seen in 6wk *Winnie* mice (females and males) was similar to age-matched controls. Analysis of CREB1 association with FOXO1 showed inhibition of CREB1-FOXO1 complex formation in male *Winnie* mice at 14wk compared to age-matched male controls (**Figure 4F**), but not in 14wk female *Winnie* mice compared to 14wk female controls (**Figure 4E**). No significant difference in CREB1-FOXO1 complex formation was seen in *Winnie* mice at 6wk (females and males).

### Winnie mice with chronic colitis exhibit increased release of GDS from the mucosa of the distal colon

To further investigate mechanisms of bone pathology at 6wk and 14wk *Winnie* mice, we examined their GDS and inflammation levels. Our results showed that GDS levels in both peak state and steady state (SS) were significantly higher in female and male *Winnie* mice at 6wk and 14wk compared to age- and sex-matched controls (**Figure 5A-D**). Evidently, *Winnie* and control mice at 14wk (females and males) showed higher levels of GDS than at 6wk.

**Figure 5:**
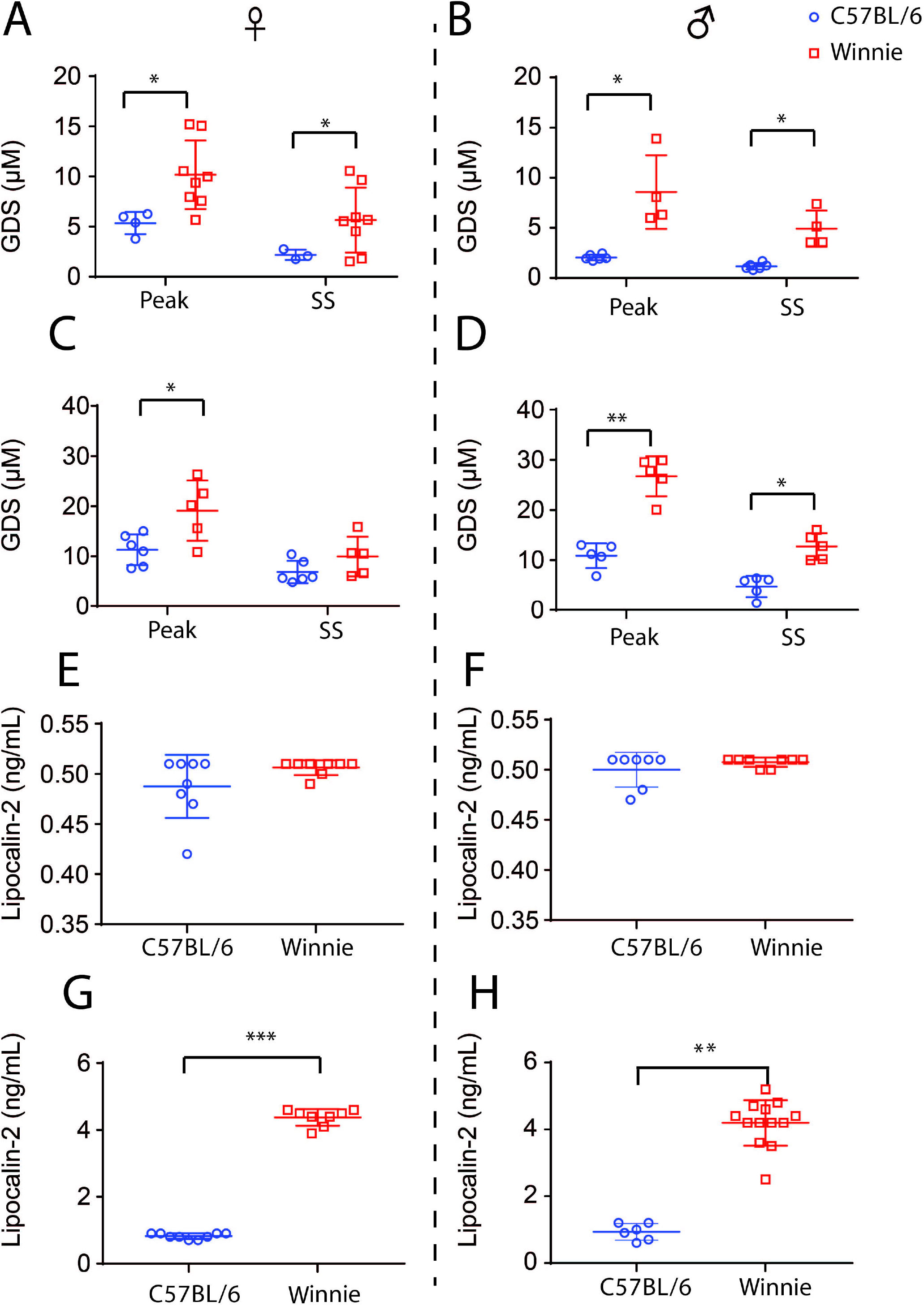
Assessment of GDS and colonic inflammation with fecal lipocalin-2. Electrochemical measurements of GDS at the mucosal surface of the colon. GDS was measured using an electrochemical technique where carbon fiber electrodes mechanically stimulate and oxidize GDS to generate a compression-evoked (peak) release of GDS which decayed back to basal levels (steady state) in both control and *Winnie* mice. (**A**) Comparison of the peak and steady state GDS levels in females *Winnie* mice and C57BL/6 and (**B**) in males at 6wk. (**C**) Comparison of the peak and steady state GDS levels in female *Winnie* mice and C57BL/6 and (**D**) in males at 14wk. Lipocalin-2 level quantified in fecal samples from females (**E**) and males (**F**) in *Winnie* mice and C57BL/6 at 6wk. (**G**) Average fecal lipocalin-2 protein levels at 14wk females and males (**H)** *Winnie* mice compared to C57BL/6. \**p*<.05, \*\**p*<.01, \*\*\**p*<.001; six replicates/sample; C57BL/6: n=4-6 animals/group, *Winnie*: n=7 animals/group.

Lipocalin-2 concentration was similar in *Winnie* mice and controls at 6wk (both females and males) (**Figure 5E, 5F)** and increased significantly in *Winnie* mice at 14wk (females and males) **(Figure 5G, 5H)** as colitis advances.

### Serum analysis for calcium, phosphorus, and vitamin D levels

Serum calcium and phosphorus levels were similar in *Winnie* mice at 6wk, 14wk and 24wk in both genders compared to age- and sex-matched controls (**Figure 6A-D)**. Vitamin D levels were not different at 6wk in *Winnie* mice compared to controls. However, vitamin D levels were significantly reduced at 14wk and 24wk in females and males in both *Winnie* and controls compared to 6wk (**Figure 6E, 6F**).

**Figure 6.**
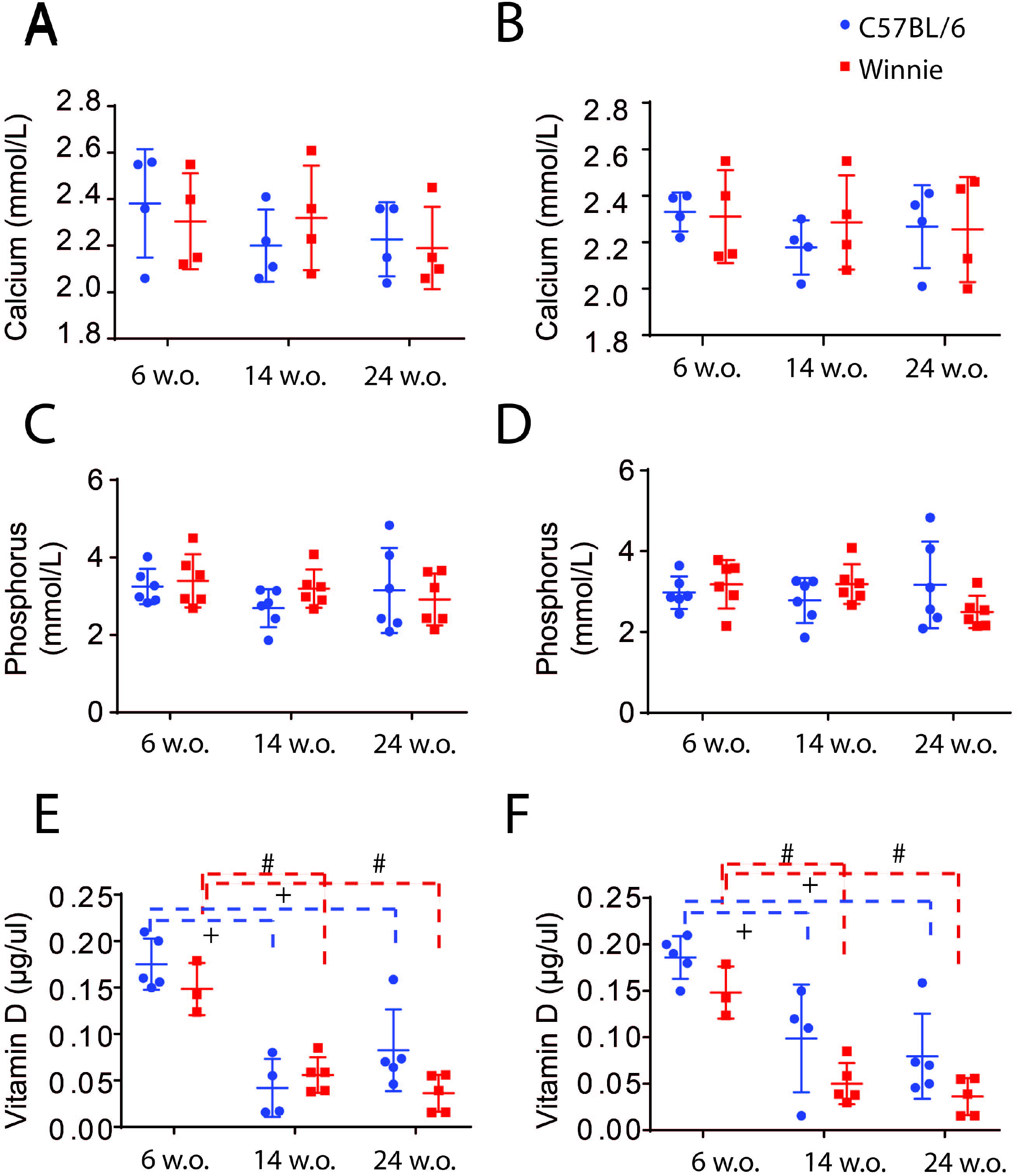
Serum levels of Calcium, phosphorus and Vitamin D in Winnie mice compared to C57BL/6 controls. **A, B)** Calcium levels in *Winnie* mice compared to C57BL/6 controls in 6wk, 14wk and 24wk in females **(A)** and males **(B)**. **C, D)** phosphorus levels in *Winnie* mice compared to C57BL/6 controls in 6wk, 14wk and 24wk in females (**C)** and males **(D)**. **E, F)** Vitamin D levels in *Winnie* mice compared to C57BL/6 controls in 6wk, 14wk and 24wk in females **(E)** and males **(F)**. n=4-10 mice/ group. Statistical significance, #*p*<.05 differences between age groups of *Winnie* mice. +*p*<.05 differences between age groups of C57BL/6 mice.

## Discussion

IBD associates with significantly high risk of developing skeletal disease, such as osteoporosis and osteopenia.^30^ However, the mechanisms underlying intestinal inflammation-associated bone loss remain elusive. In this study, we analyzed the phenotypic and functional characteristics of cortical and trabecular bone as well as the cellular and functional mechanisms underlying bone loss in a *Winnie* model of spontaneous chronic colitis. We found significant and reproducible alterations in bone structure, formation, and mechanical properties. Our results demonstrate that *Winnie* mice have lower volumetric bone mineral densities and deteriorated bone microstructure leading to fewer, thin, and more separated trabeculae, and reduced cortical thickness. To the best of our knowledge, this is the first real-time study in which the bone pathology throughout the onset and progression of chronic colitis in *Winnie* mice in both genders has been addressed. We observed skeletal deterioration with advancing age starting from the onset of colitis symptoms after 6wk, and gradually increasing from 14wk (progressive) to 24wk (severe colitis) in females and males. We have established that osteopenic phenotype in *Winnie* mice is a consequence of increased bone resorption and decreased bone formation. Paradoxically, we observed loss of some bone parameters in *Winnie* mice at 6wk without any clinical symptoms of colitis. Male *Winnie* mice at 14wk and 24wk were more susceptible to bone loss. This was evidenced by more significant deterioration of some bone parameters (trabecular thickness, elastic modulus) in male compared to female *Winnie* mice at 14wk and 24wk. Our results demonstrated elevated GDS levels starting from 6wk and progressively increasing in older mice. This might be one of the underlying causative factors of bone loss. Our study provides evidence that elevated GDS binding to its receptor on bone (5-HTR1B) initiates association/dissociation of downstream molecular regulators and contributes to decreased bone formation in colitis. Our data showed disease severity-dependent alteration of bone microarchitecture in the *Winnie* mice model of colitis. This significant alteration in bone micro-architecture is in agreement with an earlier report on a model of DSS colitis, which was conducted only in male mice and sustained for short duration of 2wk.^11^

Biomechanical studies have demonstrated that the structural properties of whole bone specimens is highly determined by the contribution of cortical bone.^31^ To understand quality and strength of cortical bone compartments, we subjected bones from *Winnie* and controls to a 3-point bending test to determine ultimate force (maximum force that breaks the bone), surrogate of intrinsic cortical bone strength and quality. The ultimate force was significantly decreased in *Winnie* mice (females and males) compared to age and sex-matched controls. We identified that age has a substantial effect on femur bone toughness (amount of energy that bone can absorb before fracture) and elastic modulus (measure of stiffness) in both genotypes.

Our study also incorporates histomorphometry, which made it possible to attribute the observed changes in bone morphology to underlying biologic processes. Significant reduction in collagen-I synthesis is present in *Winnie* mice (females and males) compared to controls. Furthermore, analysis of the cellular mechanisms of bone loss revealed that the number of osteoblasts was reduced (all ages and genders) with a concomitant elevation in osteoclasts relative to the bone volume. These results are in concordance with bone studies performed in IL-2-deficient and CD4^+^CD45RB^Hi^ T cells transfer colitis models.^32,33^ Our findings in the *Winnie* mouse model of colitis demonstrate that an increasingly complex network of interactions leads to increased bone turnover, with bone resorption exceeding formation, which accelerates the rate of bone loss concomitant to colitis. This is consistent with the earlier study performed in an acute colitis model.^11^

Furthermore, the osteoid volume and total bone volume were significantly reduced in all 3 age groups and genders compared to age- and sex-matched controls indicating low rate of bone mineralization. Consistent to this data, our *ex vivo* analysis of bone marrow showed that *Winnie* mice had low calcium deposition, which is characteristic of matrix formation.

Reduction in bone mineralization is in concordance to *in vitro* studies in which organ cultures of fetal rat parietal bones showed reduction of minerals and osteoid when treated with sera of CD patients.^34^

Our dynamic histomorphometry studies established lower rates of bone formation (BFR) and mineral apposition (MAR) (activity of the average osteoblasts, representative of new bone deposition) in *Winnie* mice compared to controls. Low BFR is consistent with earlier studies in a DSS-colitis model which demonstrate decreased BFR, but not MAR, during the acute phase of colitis^12^. However, our mouse model showed depletion in both BFR and MAR. Furthermore, our *ex vivo* bone marrow results confirmed that low BFR and MAR due to decreased differentiation ability of BM-MSCs down the osteoblast lineage in *Winnie* mice compared to controls.

The consistent feature of chronic colitis is increased GDS availability, which could have a negative impact on osteoblastogenesis in this model. Our earlier study in *Winnie* mice reports that GDS is at the maximum high level after 14wk, and ceases to rise after 24wk.^26^ High GDS levels have also been evident in TNBS and DSS-induced colitis in mice and guinea-pigs.^35^ Serotonin transporter (SERT) mRNA and protein levels were reduced in the colon of UC patients^36^ adding assurance to these findings.

Using *Winnie* mouse model of spontaneous chronic colitis, we showed for the first time that elevated GDS cross talks with molecular pathways involved in bone formation. GDS can interact with FOXO1 to regulate bone metabolism.^37^ GDS binding to serotonin receptor 5-HTR1B facilitates the proliferation of osteoblasts by maintaining the interaction of FOXO1 with CREB and ATF4, whereas an increase in GDS levels disrupts the interaction between FOXO1 and CREB and thereby inhibits the osteoblast proliferation.^37^ Consistently, our study reports that high levels of peripheral GDS in *Winnie* mice bind to 5-HTR1B on BM-MSCs to regulate transcriptional activity of FOXO1 which was also confirmed by qPCR. To elucidate downstream molecular mediators of GDS in bone, immunohistochemical analysis unveiled dissociation of FOXO1 and CREB1 complex in 14wk *Winnie* male mice. This relieves the suppressive effect of CREB on FOXO1, resulting in decreased osteoblast proliferation compared to a strong association of CREB and FOXO1 in the control group. Additionally, FOXO1 (CREB-free) association with ATF4 was observed in 14wk *Winnie* female mice, in which ATF4 promotes the transcriptional activity of FOXO1, contributing to suppression of osteoblast proliferation (**Figure 7**). In contrast, 6wk *Winnie* mice showed no significant change in association/dissociation of complexes with FOXO1 compared to 6wk controls. This is agreeable with similar 5-HTR1B expression levels in 6wk *Winnie* mice and age-matched controls.

**Figure 7.**
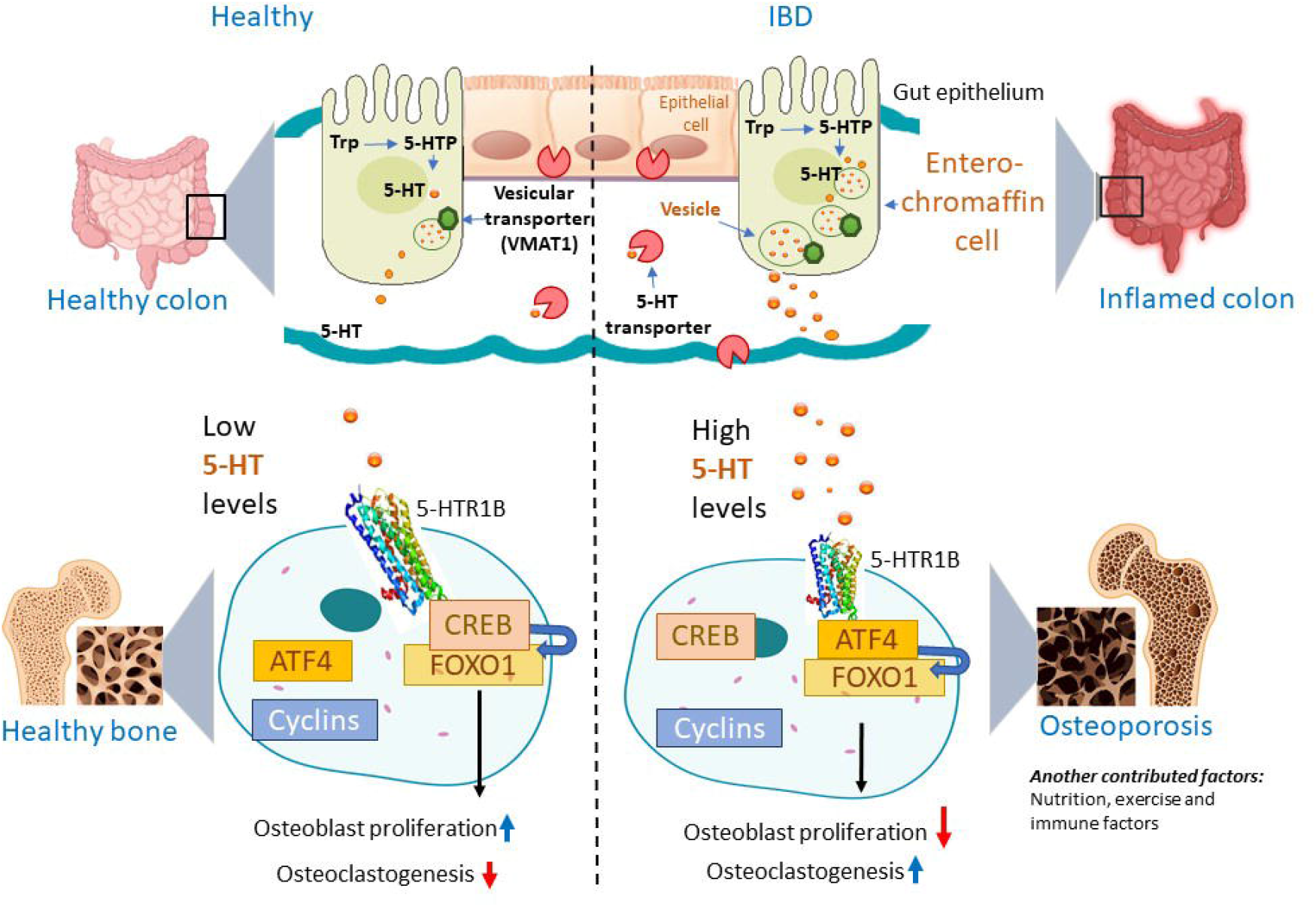
Model of mechanisms for decreased bone formation in IBD compared to healthy subjects. Gut derived serotonin (GDS) synthesis begins with its precursor amino acid L-tryptophan (Trp) converted by the rate-limiting enzyme tryptophan hydroxylase (Tph1), to 5-hydroxytryptophan (HTP). HTP is then converted by aromatic L-amino acid decarboxylase to circulating GDS and carried by vesicular transporter (VMAT1) inside the entero-chromaffin cells in the gut epithelium. Under normal physiological conditions, there are very low levels of circulating GDS to bind HTR1B receptors as most of GDS is transported intracellularly by plasmalemmal serotonin transporter and thereby gets inactivated by monoamine oxidases into 5-hydroxyindoleacetic acid. Nevertheless, under pathological conditions of colitis, GDS inhibits osteoblast proliferation as higher levels of 5HT enter the circulation because of decreased SERT expression by intestinal cells, leading to a greater activation of HTR1B receptors in bone, which has a negative impact on bone mass. CREB1 is necessary for optimal expression of cyclin D1 and for normal osteoblast proliferation. HTR1B signaling inhibits both expression and phosphorylation dependent activation of the transcription factor CREB.

These findings identify FOXO1 as the molecular node of an intricate transcriptional machinery that confers the signaling of high GDS levels to inhibit bone formation. These data indicate that, in high GDS levels, there is a shift in FOXO1 target genes from CREB- to ATF4-dependent responses. As there is no direct proportion of increment of dissociation of CREB-FOXO1 and association of ATF4-FOXO1 complex, we can implicate that GDS-activated FOXO1 may interact in BM-MSCs with a set of proteins distinct from those used under physiological conditions, especially in 14wk *Winnie* mice, where CREB-free FOXO1 is more available but ATF4-FOXO1 association is not significant compared to 14wk controls. Our findings indicate that the interaction of FOXO1 with ATF4 or CREB under the control of GDS can alter the outcome on osteoblast proliferation by shifting to ATF4-dependent responses. In conclusion, findings reported here strongly support the hypothesis that changes in GDS signaling can contribute to the bone loss associated with colitis. Consistently, Lavoie et al (2019) demonstrated deleterious effect on bone caused by high GDS levels in the DSS model of colitis.^25^

It has been speculated that GDS triggers immune cells for the production of pro-inflammatory mediators in the context of intestinal inflammation and inflammatory joint diseases.^24,38^ Likewise, GDS may have a pro-inflammatory role in causing bone loss in colitis. Inflammation-independent studies conducted in knock-out mouse models (LRP5^-/-^, FOXO1^-/-^) also revealed elevated GDS as a causative agent for low bone phenotypes. Therefore, our study provides evidence that bone loss associated with colitis involves the anti-anabolic effect of GDS, at least in part, along with inflammation.

We observed changes in few bone parameters (osteoblast number, collagen-1, total collagen) in 6wk *Winnie* mice compared to age-matched controls, although no clinical symptoms of colitis were present at this age consistent with previous studies.^15^ Electrochemical measurements revealed elevated GDS levels at 6wk in *Winnie* mice; qRT-PCR analysis showed non-significant increase in 5-HTR1B levels at 6wk (females and males) *Winnie* mice compared to controls suggesting a non 5-HTR1B dependent mechanism.^39^ These findings suggest that bone loss observed in 6wk *Winnie* mice might be attributed to elevated GDS levels. Unlike 6wk *Winnie* mice, significantly higher levels of inflammation correlated positively with severe bone loss at 14wk and 24wk.

We also explored the relative contribution of malabsorption to bone mass and structure. Similar serum levels of calcium and phosphorus in control and *Winnie* mice in all age groups (females and males) rule out the possibility of malabsorption of these minerals. Similarly, studies in humans have concluded that nutritional factors, vitamin deficiency, and malabsorption are not major contributing factors to bone mineral density loss.^40,41^ Also, multiple animal models of colitis, including DSS, HLA-B27 transgenic rats and IL10^-/-^ knockout mice display reduced bone mass, bone volume fraction and bone strength,^11,12^ and the effects cannot be explained solely by impaired nutritional absorption.^42^ Consistently, we also found in this study that all mice consumed, on average, the same amount of food or water over the course of the study (our unpublished observations), and hence decreased food intake was not a causative agent.

## Conclusion

Taken together, our results indicate that the onset and severity of intestinal inflammation parallels the bone loss phenotype, characterized by decreased cortical and trabecular bone morphometry, osteoid, osteoblast numbers, collagen-I, and dynamic measures of bone formation. Bone loss in *Winnie* mice is a consequence of elevated bone turnover, with a higher rate of osteoclastogenesis than osteoblastic bone formation. Bone loss affects both female and male *Winnie* mice although the magnitude of bone loss differs by gender. Higher skeletal deterioration in male *Winnie* mice is concordant to higher GDS levels. Collectively, our findings support the hypothesis that *Winnie* mice are osteopenic and can be used to explore the mechanisms and pharmacological interventions for bone loss in IBD. Our findings support the concept that elevated GDS can have a negative impact on bone growth concomitant to colitis. A potential pitfall of our study is not demonstrating IBD-related bone effects after blocking serotonin. Therefore, future studies are warranted to investigate the relative roles of therapeutics to mitigate the risk of this extraintestinal manifestation of IBD.

## References

1. Kaplan GG, Ng SC. Understanding and Preventing the Global Increase of Inflammatory Bowel Disease. Gastroenterology 2017;152:313–321 e2.

2. Vedamurthy A, Ananthakrishnan AN. Influence of Environmental Factors in the Development and Outcomes of Inflammatory Bowel Disease. Gastroenterol Hepatol (N Y) 2019;15:72–82.

3. Knights D, Lassen KG, Xavier RJ. Advances in inflammatory bowel disease pathogenesis: linking host genetics and the microbiome. Gut 2013;62:1505–10.

4. Bourikas LA, Papadakis KA. Musculoskeletal manifestations of inflammatory bowel disease. Inflamm Bowel Dis 2009;15:1915–24.

5. Vazquez MA, Lopez E, Montoya MJ, et al. Vertebral fractures in patients with inflammatory bowel disease compared with a healthy population: a prospective casecontrol study. BMC Gastroenterol 2012;12:47.

6. Bartko J, Reichardt B, Kocijan R, et al. Inflammatory Bowel Disease: A Nationwide Study of Hip Fracture and Mortality Risk after Hip Fracture. J Crohns Colitis 2020.

7. van Staa TP, Cooper C, Brusse LS, et al. Inflammatory bowel disease and the risk of fracture. Gastroenterology 2003;125:1591–7.

8. Ghishan FK, Kiela PR. Advances in the understanding of mineral and bone metabolism in inflammatory bowel diseases. Am J Physiol Gastrointest Liver Physiol 2011;300:G191–201.

9. Katz S, Weinerman S. Osteoporosis and gastrointestinal disease. Gastroenterol Hepatol (N Y) 2010;6:506–17.

10. Sgambato D, Gimigliano F, De Musis C, et al. Bone alterations in inflammatory bowel diseases. World journal of clinical cases 2019;7:1908–1925.

11. Hamdani G, Gabet Y, Rachmilewitz D, et al. Dextran sodium sulfate-induced colitis causes rapid bone loss in mice. Bone 2008;43:945–950.

12. Harris L, Senagore P, Young VB, et al. Inflammatory bowel disease causes reversible suppression of osteoblast and chondrocyte function in mice. American journal of physiology. Gastrointestinal and liver physiology 2009;296:G1020–G1029.

13. Irwin R, Raehtz S, Parameswaran N, et al. Intestinal inflammation without weight loss decreases bone density and growth. Am J Physiol Regul Integr Comp Physiol 2016;311:R1149–R1157.

14. Irwin R, Lee T, Young VB, et al. Colitis-induced bone loss is gender dependent and associated with increased inflammation. Inflammatory bowel diseases 2013;19:1586–1597.

15. Heazlewood CK, Cook MC, Eri R, et al. Aberrant mucin assembly in mice causes endoplasmic reticulum stress and spontaneous inflammation resembling ulcerative colitis. PLoS Medicine 2008;5:0440–0460.

16. Eri RD, Adams RJ, Tran TV, et al. An intestinal epithelial defect conferring ER stress results in inflammation involving both innate and adaptive immunity. Mucosal Immunology 2010;4:354–364.

17. Buisine MP, Desreumaux P, Leteurtre E, et al. Mucin gene expression in intestinal epithelial cells in Crohn’s disease. Gut 2001;49:544–51.

18. Van Klinken BJ, Van der Wal JW, Einerhand AW, et al. Sulphation and secretion of the predominant secretory human colonic mucin MUC2 in ulcerative colitis. Gut 1999;44:387–93.

19. Robinson AM, Rahman AA, Carbone SE, et al. Alterations of colonic function in the Winnie mouse model of spontaneous chronic colitis. American Journal of Physiology-Gastrointestinal and Liver Physiology 2016;312:G85–G102.

20. Rahman AA, Robinson AM, Jovanovska V, et al. Alterations in the distal colon innervation in Winnie mouse model of spontaneous chronic colitis. Cell and Tissue Research 2015;362:497–512.

21. Robinson AM, Gondalia SV, Karpe AV, et al. Fecal Microbiota and Metabolome in a Mouse Model of Spontaneous Chronic Colitis: Relevance to Human Inflammatory Bowel Disease. Inflamm Bowel Dis 2016;22:2767–2787.

22. Al Saedi A, Sharma S, Summers MA, et al. The multiple faces of tryptophan in bone biology. Exp Gerontol 2020;129:110778.

23. Duque GA-O, Vidal C, Li W, et al. Picolinic acid, a catabolite of tryptophan, has an anabolic effect on bone in vivo. LID −10.1002/jbmr.4125 [doi]. J Bone Miner Res 2020.

24. Yadav VK, Ryu J-h, Suda N, et al. Lrp5 controls bone formation by inhibiting seotonin synthesis in the duodenum: an entero-bone endocrine axis. Cell 2008;135:825–837.

25. Lavoie B, Roberts JA, Haag MM, et al. Gut-derived serotonin contributes to bone deficits in colitis. Pharmacological Research 2019;140:75–84.

26. Stavely R, Fraser S, Sharma S, et al. The Onset and Progression of Chronic Colitis Parallels Increased Mucosal Serotonin Release via Enterochromaffin Cell Hyperplasia and Downregulation of the Serotonin Reuptake Transporter. Inflammatory Bowel Diseases 2018;24:1021–1034.

27. Bouxsein ML, Boyd SK, Christiansen BA, et al. Guidelines for assessment of bone microstructure in rodents using micro-computed tomography. J Bone Miner Res 2010;25:1468–86.

28. McGregor NE, Murat M, Elango J, et al. IL-6 exhibits both cis- and trans-signaling in osteocytes and osteoblasts, but only trans-signaling promotes bone formation and osteoclastogenesis. J Biol Chem 2019;294:7850–7863.

29. Yang R, Chen J, Zhang J, et al. 1,25-Dihydroxyvitamin D protects against age-related osteoporosis by a novel VDR-Ezh2-p16 signal axis. Aging Cell 2020;19:e13095.

30. Laakso S, Valta H, Verkasalo M, et al. Compromised peak bone mass in patients with inflammatory bowel disease--a prospective study. J Pediatr 2014;164:1436–43 e1.

31. Burrows M, Liu D, Moore S, et al. Bone microstructure at the distal tibia provides a strength advantage to males in late puberty: An HR-pQCT study. Journal of Bone and Mineral Research 2010;25:1423–1432.

32. Ashcroft AJ, Cruickshank SM, Croucher PI, et al. Colonic Dendritic Cells, Intestinal Inflammation, and T Cell-Mediated Bone Destruction Are Modulated by Recombinant Osteoprotegerin. Immunity 2003;19:849–861.

33. Byrne FR, Morony S, Warmington K, et al. CD4+CD45RBHi T cell transfer induced colitis in mice is accompanied by osteopenia which is treatable with recombinant human osteoprotegerin. Gut 2005;54:78–86.

34. Hyams JS, Wyzga N, Kreutzer DL, et al. Alterations in bone metabolism in children with inflammatory bowel disease: an in vitro study. J Pediatr Gastroenterol Nutr 1997;24:289–95.

35. Linden DR, Chen J-X, Gershon MD, et al. Serotonin availability is increased in mucosa of guinea pigs with TNBS-induced colitis. American Journal of Physiology-Gastrointestinal and Liver Physiology 2015;285:G207–G216.

36. Coates MD, Mahoney CR, Linden DR, et al. Molecular defects in mucosal serotonin content and decreased serotonin reuptake transporter in ulcerative colitis and irritable bowel syndrome. Gastroenterology 2004;126:1657–1664.

37. Kode A, Mosialou I, Silva BC, et al. FOXO1 orchestrates the bone-suppressing function of gut-derived serotonin. Journal of Clinical Investigation 2012;122:3490–3503.

38. Ragab O, Khairy N, Taha R, et al. Serum serotonin in rheumatoid arthritis patients: Relation to rheumatoid factor positivity, clinical manifestations and fibromyalgia. The Egyptian Rheumatologist 2018;40:149–153.

39. de Vernejoul M-C, Collet C, Chabbi-Achengli Y. Serotonin: good or bad for bone. BoneKEy Reports 2012;1:1–6.

40. Haderslev KV, Tjellesen L, Sorensen HA, et al. Alendronate increases lumbar spine bone mineral density in patients with Crohn’s disease. Gastroenterology 2000;119:639–646.

41. Jahnsen J, Falch JA, Mowinckel P, et al. Vitamin D status, parathyroid hormone and bone mineral density in patients with inflammatory bowel disease. Scandinavian journal of gastroenterology 2002;37:192–199.

42. Ghishan FK, Kiela PR. Advances in the understanding of mineral and bone metabolism in inflammatory bowel diseases. American journal of physiology. Gastrointestinal and liver physiology 2011;300:G191–G201.

