## Supplementary Methods for "Skeletal Phenotype and Mechanisms of Bone Loss in *Winnie* Mice as a Model for Inflammatory Bowel Disease"

**Materials and Methods (Detailed):**

*Micro-CT analysis*

Structural analysis of the bone was performed using micro computed tomography (*Micro*-CT) in the left femur after removal of soft tissue and overnight fixation in 4% paraformaldehyde. The distal metaphysis was analyzed with a Micro-CT instrument (Skyscan, 1272, Bruker, Belgium )). Regions of interest (ROIs) in the distal portion of each femur started from the proximal border of the growth plate (0.0–2.5 mm from the growth plate border). Image acquisition was performed at 50 kV and 200 μA, with a 0.4° rotation between frames. The two-dimensional images were used to generate three-dimensional reconstructions to obtain quantitative data with the 3D Creator manufacturer’s software. Nomenclature and abbreviations of 3D Micro-CT parameters follow the recommendations of the American Society for Bone and Mineral Research. The following data were obtained: bone volume to tissue volume ratio (BV/TV), trabecular thickness (Tb.Th), trabecular number (Tb.N) and trabecular separation (Tb.Sp).

*Three-point bending test*

Three-point bending tests were conducted as described previously^1^ with a Biodynamic 5500 test instrument (TA Instruments, New Castle, DE). Male and female bones were tested with a constant span of 9 cm. Before testing, tibiae were kept moist in gauze swabs soaked in PBS. The bone was positioned horizontally with the anterior surface upward, centred on the supports, and the pressing force was directed vertically to the mid-shaft of the bone. Each bone was compressed at a constant 0.5 mm/s until failure. WinTest software was used to collect the load-displacement data at 250 data points per second for a total of 10 s. Structural properties, including ultimate force, yield force, stiffness, and energy to failure endured by the tibiae, were calculated from load/displacement data.^2^ The yield point was determined from the load deformation curve at the point at which the curve deviated from linear. Widths of the cortical mid-shaft in the mediolateral and anteroposterior directions, moment of inertia, and average cortical thickness determined by Micro-CT were combined with three-point bending data to calculate material-specific properties.^3^

*Static Histomorphometry*

The details of these methods were described previously.^4^ PFA fixed tibiae bones were decalcified in 14% EDTA (pH 7.4) for 2 wks. Bones were embedded in paraffin and 5 µm sections were cut using a rotary microtome. Sections were then stained with H&E for osteoblasts and tartrate-resistant acid phosphatase (TRAP) for osteoclast staining using standard protocol (ref). Naphtol-AS-TR (Sigma-Aldrich, Australia) was used as substrate for both enzymes; Fast Blue BB salt (Sigma-Aldrich, Australia) was used as a coupler for alkaline phosphatase. Von Kossa staining was used to perform histomorphometry.

*Total-collagen staining*

Tissue sections were deparaffinized and hydrated through xylenes and graded alcohol series. Subsequently, they were incubated in picronitric acid-direct red solution for 0.5-1 hours at RT and washed. This was followed by counterstaining with hematoxylin, then rinsing in running water. Dehydrated and clear tissue sections through graded alcohol series and xylenes were mounted with Kaiser's glycerol jelly.

*Immunostaining for type I collagen*

Sections were de-waxed and rehydrated with phosphate-buffered saline (PBS), then incubated in 0.5% hyaluronidase at 37°C for 30 min, and washed with 3xPBS. This was followed by incubation in 3% hydrogen peroxide for 30min, washing and incubation in 10% Rabbit serum (with 0.5% BSA-PBS). Subsequently, sections were stained with goat anti-type 1 collagen (1:100) overnight at RT and washed. The sections were stained with Rabbit anti-goat IgG (1:100) at 28°C for 1hr then washed. Subsequently, they were stained with Elite-ABC at 28°C for 1 hr (Vector Elite ABC kit).^4^ Finally, staining was done with DAB followed by counterstaining with haematoxylin.

*Plastic-embedded sectioning*

Tibiae bones were incubated with 100% ethanol for 24 hrs, followed by a second incubation with 95% ethanol for 24 hrs, a third incubation with 90% ethanol for 24 hrs, and a final incubation with 70% ethanol at 4°C. Bones were rinsed with PBS and embedded in polymethyl methacrylate (MMA). Serial 4- to 6- µm sections of MMA-embedded tissues were then stained with ALP for osteoblast and toluidine blue for osteocytes, Von Kossa histochemical staining for bone matrix and tartrate resistance acid phosphatase activity for osteoclasts.^4^ The primary histomorphometric data was obtained using Bioquant image analysis software.

*Dynamic Histomorphometry*

The fluorochrome, calcein was injected (10mg/kg dissolved in 2.0% sodium bicarbonate, pH 7) at 0.1ml per 10g body weight at 7 and 2 days prior to bone/muscle harvest from the mice. Both side tibiae and femora were dissected out for analysis from 6wk, 14wk and 24wk of *Winnie* and C57BL/6 mice. One femur from each animal in each group was removed at the time of euthanasia, fixed in 70% ethanol, dehydrated, and embedded undecalcified in methylmethacrylate (J-T Baker, Phillipsburg, NJ, USA). At 50 μm intervals, longitudinal sections of 5 and 8 μm thick were cut using a polycut-E microtome (Reichert-Jung Leica, Heerbrugg, Switzerland), placed on gelatin-coated glass slides, deplastified and stained with Goldner's trichrome. Histomorphometry was performed with a semi-automatic image analyzing system combining a microscope equipped with a camera lucida and digitizing tablet linked to a computer using the Bioquant Software^®^. Nomenclature and abbreviations of histomorphometry parameters follow the recommendations of the American Society for Bone and Mineral Research.^5^

*Ex vivo bone marrow cultures and staining*

Bone marrow (BM) flushing was done within 1-2 hrs after harvesting tibia and collected in fresh media (alpha-MEM, 10% FBS, 1% antibiotics-antimycotics, 1% Glutamax, 10% FBS). BM Cells were incubated at 37°C with 5% humidified CO_2_. Bone marrow stromal cells BMSCs were isolated by their adherence to tissue culture plastic. Medium was aspirated and replaced with fresh medium every 2 to 3 days to remove non-adherent cells.

To study the osteogenic differentiation potential of BMSCs, total of 0.1 million cells were diluted in osteogenic medium (prepared with alpha-MEM (Sigma), 10% FBS, 0.2 mM dexamethasone, 10 mmol/L β-glycerol phosphate and 50 μg/ml ascorbic acid) and plated in 24-well plates at passage 2. Media was aspirated and replaced with fresh osteogenic medium every 3 days. After 7 days in culture, cells were washed with 1X PBS, fixed in 10% formalin for 2min, again washed with 1X PBS, and then stained for alkaline phosphatase using SIGMAFAST™ p-Nitrophenyl phosphate tablets (N1891) according to the manufacturer's instructions. The colonies with more than 10% of cells staining positive for alkaline phosphatase (ALP) were considered as colony-forming units–osteoblasts (CFU-OBs).

Alizarin red (Merck, Millipore) staining was performed to assess mineralized bone nodule formation. After 14 days of culture of BM-MSCs in osteogenic medium, cells were fixed in 70% ice-cold ethanol for 1hr and rinsed with distilled water. Cells were stained with 40 mM Alizarin Red S (pH 4.2) for 10 min with gentle agitation. The level of Alizarin Red S staining was observed under light microscopy. For quantification of mineralization, 10% cetylpyridinium chloride (CPC) (prepared in 10 mM sodium phosphate at pH 7.0) were added to stained wells for 2 hr at RT to de-stain. Alizarin Red staining was then quantified by measuring the absorbance of the eluted stain at 562nm using a spectrophotometer.

*Double Immunofluorescence labeling*

Freshly isolated BM was flushed out using 1cc syringe in alpha-MEM (Sigma), 10%FBS, 10% antibiotics/antimycotics, 1% glutamax. MSCs were cultured till passage 2. MSCs were trypsinized using triplex (Invitrogen), pelleted and seeded in chamber slides (Nunc). After 48hrs, MSCs were covered to a depth of 2–3 mm for fixation using 4% formaldehyde diluted in 1X PBS for 15 min at room temperature. Fixative was aspirated and rinsed three times in 1X PBS for 5 min each. Fixed cells were blocked in blocking buffer for 60 min (1X PBS / 5% normal donkey serum / 0.3% Triton™ X-100) to block non-specific Ab binding and for permeabilization. Blocking solution was aspirated and then cells were incubated with primary Abs, mouse monoclonal FOXO1 (14952, CST) and rabbit polyclonal anti-phospho CREB1 conjugate (06-519-AF647, Millipore Sigma) simultaneously in Ab dilution buffer (1X PBS / 1% BSA / 0.3% Triton™ X-100) for overnight at 4°C in dark in a humified chamber. Similarly, other wells of MSCs were incubated with primary Abs, mouse monoclonal FOXO1 and rabbit polyclonal ATF4 (ab23760, Abcam) simultaneously and washed with PBS. Both types of primary Ab labelled MSCs (CREB1:FOXO1) and (ATF4: FOXO1) were then incubated with secondary antibodies, FITC-conjugated anti-mouse IgG (1:250) and Cy3-conjugated rabbit anti-mouse IgG (1:200) for 2 hrs at RT in dark. Confocal immunofluorescence images were acquired. Labelled MSCs were visualized using an Eclipse Ti confocal microscope (Nikon, Tokyo, Japan) using 60X objective.

*Serum analysis*

The serum sample were prepared by centrifugation of the blood collected by cardiac puncture (1.013 g for 15 min 4 °C), then stored at −80°C for biochemical assay. Serum calcium, phosphorus and vitamin D levels were assayed by ASAP Laboratory services, Melbourne, Australia according to the manufacturer’s instructions. This calcium procedure is based on calcium ions (Ca^2+^) reacting with Arsenazo III (2,2’-[1,8-Dihydroxy-3,6-disulphonaphthylene-2,7-bisazo]- bisbenzenesonic acid) to form an intense purple-colored complex. Magnesium does not significantly interfere in calcium determination using Arsenazo III. In this method, the absorbance of the Ca-Arsenazo III complex is measured dichromatically at 660/700 nm using spectrophotometer. The resulting increase in absorbance of the reaction mixture is directly proportional to the calcium concentration in the sample. Phosphorus measured based on a modification of the method developed by Daly and Ertingshausen in which inorganic phosphate reacts with molybdate to form a heteropolyacid complex. The use of a surfactant eliminates the need to prepare a protein free filtrate. The absorbance at 340/380 nm is directly proportional to the inorganic Phosphorus level in the sample. Similarly, vitamin D levels were measured by radioimmunoassay (Diagnostic Products).

*Gut-derived serotonin measurement*

GDS measurements were done using an electrochemical technique^6-10^. Segments of the distal colon were cut along the mesenteric border under a dissecting microscope and loosely pinned mucosal side up in a silicon-lined recording chamber. The chamber was super-fused with carbogen (95% O_2_ and 5% CO_2_) bubbled physiological Krebs solution (composition in mmol/L: NaCl, 117; NaH_2_PO4, 1.2; MgSO4, 1.2; CaCl_2_, 2.5; KCl, 4.7; NaHCO_3_, 25; and glucose, 11) at 35°C at a flow rate of ~5mL/minute. Tissues were equilibrated for 60 min before amperometric recordings of GDS oxidation commenced. Microelectrodes were prepared by insulating a 7μm carbon fiber with a borosilicate glass capillary (outer diameter, 1.5 mm; inner diameter, 0.86 mm; Harvard Apparatus, Holliston, MA, USA) leaving ∼200 μm of carbon fiber exposed at the recording tip. Within the capillary, a pellet of woods metal was used to join the remaining carbon fiber and copper wire to provide a connection point for the head-stage. Carbon fiber electrodes were voltage clamped at +400 mV; GDS oxidation was detected as a positive current deflection. Recordings of the current generated by the oxidation of GDS were made using a VA-10 amplifier (NPI Electronics, Tamm, Germany), digitized at 1–5 kHz (Digidata 1440; Axon Instruments, Union City, CA, USA) to a personal computer using PClamp 9.0 (MDS Analytical Technologies, Mississauga, ON, Canada) with 0.5 kHz filtering with a 50 Hz notch filter). All manufactured electrodes were individually calibrated with a 10μL spritz of 10μM serotonin hydrochloride (Sigma-Aldrich, Sydney, Australia) in Krebs solution before performing recordings. A precision micromanipulator was used to compress the mucosa with the carbon fiber microelectrode to induce mechanically stimulated GDS release (peak) and the decay of GDS back to baseline levels (steady state).

*Assessment of Inflammation by Lipocalin-2 assay*

To assess the level of colonic inflammation in *Winnie* mice, fecal lipocalin (Lcn)-2 was measured using Abcam kit (ab119601, Mouse lipocalin-2 ELISA kit, NGAL). Lcn-2 was quantified by ELISA, as previously described.^2^ Mice fecal pellets were collected and snap frozen at LQN, and were then transferred to -80^o^C for the long-term. Frozen fecal samples were reconstituted in 50µl of PBS containing 0.1% Tween 20 (100 mg/ml) and vortexed for 20 min to get a homogenous fecal suspension. These samples were then centrifuged for 10 min at 12,000 rpm and 4°C. Clear supernatants were collected and stored at −20°C until analysis. On the day of experiment, fecal samples were thawed for 2 hrs at RT and all materials and prepared reagents were equilibrated to RT prior to use. Serially diluted standards were prepared immediately prior to use. Lipocalin-2 mouse lyophilized recombinant protein standard sample was reconstituted the by adding 1 mL Sample Diluent NS by pipette. It was mixed thoroughly and gently and kept for 10 minutes. This is the 3333 pg/mL Stock Standard Solution. 273 μL Sample Diluent NS was added into tube number 1 and 150 μL of Sample Diluent NS into numbers 2-8. Number 8 contains no protein and is the Blank control. 50 μL of all samples or standards were added to appropriate wells. 50 μL of the Antibody Cocktail (10X Capture Antibody and 300 μL 10X Detector Antibody with 2.4 mL Antibody Diluent 5B) was added to each well. The plate was sealed and incubated for 1 hour at room temperature on a plate shaker set to 400 rpm. Each well was washed with 3 x 350 μL 1X Wash Buffer PT. Washing by aspirating or decanting from wells was done three times. After the last wash, the plate was inverted and blotted against clean paper towels to remove excess liquid. 100 μL of TMB Development Solution was added to each well and incubated for 10 minutes in the dark on a plate shaker set to 400 rpm. Lastly, 100 μL of Stop Solution was added to each well and then placed on a plate shaker for 1 minute to mix. The OD was recorded at 450 nm.

*Quantitative Real-time PCR:*

Purified total RNA was extracted from fresh bone marrow derived from mesenchymal stem cells at passage 2 using Qiagen Minikit (Qiagen). RNA was treated with Genomic DNA extraction columns (Qiagen) to remove residual genomic DNA. RNA quantity (Absorbance at 260) and purity (A260/A280nm) was assessed with a nanodrop spectrophotometer. RNA with absorbance ratio (260/280nm) ratio of 2.1 only were included in the study. Total DNase-treated RNA (500 ng from n = 3 animals/group) was denatured for 5 mins at 65°C and reverse transcribed for 1 hour at 50°C using Superscript III Reverse Transcriptase Super-Mix (Thermo Fisher Scientific, Australia). The mRNA levels of HTR1B, FOXO1, and reference gene GAPDH were detected using PrimePCR™ Assays (Bio-Rad), which comprised predesigned and validated primer pairs specific to each gene. Amplification conditions were 95°C for a 2-minute cycle followed by 40 cycles at 95°C for 15 seconds and 60°C for 1 min. RT-qPCR was performed in a Bio-Rad CFX96 real-time thermal cycler in a reaction volume of 20 µL consisting of 10 µL 2×SsoAdvanced™ SYBR® Green supermix (Bio-Rad), 4  µL diluted cDNA template, 1 µL gene-specific primer pair (PrimePCR™ Assay, BioRad), and 5 µL nuclease-free water. The data was processed using the Bio-Rad CFX Manager™ software (version 3.1), using a constant threshold level to determine crossing point (Ct) values.

**References**

1.	**Cho DC,** Brennan HJ, Johnson RW, et al. Bone corticalization requires local SOCS3 activity and is promoted by androgen action via interleukin-6. Nat Commun 2017;8:806.

2.	**Jepsen KJ**, Silva MJ, Vashishth D, et al. Establishing biomechanical mechanisms in mouse models: practical guidelines for systematically evaluating phenotypic changes in the diaphyses of long bones. J Bone Miner Res 2015;30:951-66.

3.	**Turner CH**, Burr DB. Basic biomechanical measurements of bone: a tutorial. Bone 1993;14:595-608.

4.	**Yang R**, Chen J, Zhang J, et al. 1,25-Dihydroxyvitamin D protects against age-related osteoporosis by a novel VDR-Ezh2-p16 signal axis. Aging Cell 2020;19:e13095.

5.	**Bouxsein ML**, Boyd SK, Christiansen BA, et al. Guidelines for assessment of bone microstructure in rodents using micro-computed tomography. J Bone Miner Res 2010;25:1468-86.

6.	**Bertrand PP**, Barajas-Espinosa A, Neshat S, et al. Analysis of real-time serotonin (5-HT) availability during experimental colitis in mouse. Am J Physiol Gastrointest Liver Physiol 2010;298:G446-55.

7.	**Bertrand PP**, Hu X, Mach J, et al. Serotonin (5-HT) release and uptake measured by real-time electrochemical techniques in the rat ileum. Am J Physiol Gastrointest Liver Physiol 2008;295:G1228-36.

8.	**Bertrand PP**. Real-time detection of serotonin release from enterochromaffin cells of the guinea-pig ileum. Neurogastroenterol Motil 2004;16:511-4.

9.	**Patel BA**. Electroanalytical approaches to study signaling mechanisms in the gastrointestinal tract. Neurogastroenterol Motil 2011;23:595-605.

10.	**Stavely R**, Robinson AM, Miller S, et al. Allogeneic guinea pig mesenchymal stem cells ameliorate neurological changes in experimental colitis. Stem Cell Res Ther 2015;6:263.
